# Evolution of the Codling Moth Pheromone Through the Member of an Ancient Desaturase Expansion

**DOI:** 10.1101/2020.12.03.410647

**Authors:** Jean-Marc Lassance, Bao-Jian Ding, Christer Löfstedt

**Author notes:** These authors contributed equally to this work (Jean-Marc Lassance, Bao-Jian Ding).

## Abstract

Defining the origin of genetic novelty is central to our understanding of the evolution of novel traits. Diversification among fatty acid desaturase (FAD) genes has played a fundamental role in the introduction of structural variation in fatty acyl derivatives. Because of its central role in generating diversity in insect semiochemicals, the FAD gene family has become a model to study how gene family expansions can contribute to the evolution of lineage-specific innovations. Here we used the codling moth (*Cydia pomonella*) as a study system to decipher the proximate mechanism underlying the production of the Δ8Δ10 signature structure of Olethreutine moths. Biosynthesis of the codling moth sex pheromone, (*E*8,*E*10)-dodecadienol (codlemone), involves two consecutive desaturation steps, the first of which is unusual in that it generates an *E*9 unsaturation. The second step is also atypical: it generates a conjugated diene system from the *E*9 monoene C_12_ intermediate via 1,4-desaturation. Here we describe the characterization of the FAD gene acting in codlemone biosynthesis. We identify 27 FAD genes corresponding to the various functional classes identified in Insects and Lepidoptera. These genes are distributed across the *C. pomonella* genome in tandem arrays or isolated genes, indicating that the FAD repertoire consists of both ancient and recent duplications and expansions. Using transcriptomics, we show large divergence in expression domains: some genes appear ubiquitously expressed across tissue and developmental stages; others appear more restricted in their expression pattern. Functional assays using heterologous expression systems reveal that one gene, Cpo_CPRQ, which is prominently and exclusively expressed in the female pheromone gland, encodes an FAD that possesses both *E*9 and Δ8Δ10-desaturation activities. Phylogenetically, Cpo_CPRQ clusters within the Lepidoptera-specific Δ10/Δ11 clade of FADs, a classic reservoir of unusual desaturase activities in moths. Our integrative approach shows that the evolution of the signature pheromone structure of Olethreutine moths relied on a gene belonging to an ancient gene expansion. Members of other expanded FAD subfamilies do not appear to play a role in chemical communication. This advises for caution when postulating the consequences of lineage-specific expansions based on genomics alone.

## Introduction

Establishing the origin of genetic novelty and innovation is central to our understanding of the evolution of novel traits. While genes can evolve *de novo* in the genome, the most common mechanism involves duplications of existing genes and the subsequent evolution of novel properties harbored by the encoded gene products. Expansions provide opportunities for specialization and evolution of novel biological functions within a lineage, including the breadth of expression via modifications of promoter architecture (Lespinet et al. 2002). The size of gene families is influenced by both stochastic processes and selection, and particularly large differences in genetic makeup can be indicative of lineage-specific adaptation and potentially associate with traits contributing to phenotypical differentiation between groups (Hahn et al. 2005, 2007). However, the precise functional consequences of such amplifications remain frequently unclear, even if expansions show readily discernible patterns. Therefore, deciphering the underlying genetic and molecular architecture of new phenotypic characters is necessary for a complete understanding of the role played by the accumulation of genetic variation through gene duplication.

For organisms relying on chemical communication, evolution of the ability to produce and detect a new type of molecule could allow for the expansion of the breadth of available communication channels and provide a medium with no or limited interference from other broadcasters. Since the identification of bombykol by Butenandt and co-workers in 1959 (Butenandt et al. 1959), a countless number of studies have contributed to revealing the diversity of fatty acid derivatives that play a pivotal role in the chemical communication of insects. As a consequence of homologies with well-studied metabolic pathways, our understanding of the molecular basis of pheromone biosynthesis from fatty acid intermediates has greatly advanced over the past two decades, highlighting the role of several multigene families (Löfstedt et al. 2016). Insect semiochemicals are synthesized in specialized cells in which fatty acyl intermediates are converted in a stepwise fashion by a combination of desaturation, chain-shortening and chain-elongation reactions followed by modifications of the carbonyl group, to cite a few possible steps (Tillman et al. 1999; Löfstedt et al. 2016 and references therein). Desaturation appears particularly important and contributes to producing the great diversity of structures observed in insect pheromones. This derives from the properties of the enzymes that are central to many uncanonical fatty acid synthesis pathways seen in insects. These enzymes can exhibit diverse substrate preference, introduce desaturation in either or both *cis* (*Z*) and *trans* (*E*) geometry, and give rise to variation in chain-length double bond position, number and configuration.

In insects, the group of proteins responsible for catalyzing desaturation reactions are fatty acyl-CoA desaturases (FADs). These membrane-bound acyl-lipid desaturases are biochemically and structurally homologous to the desaturases ubiquitously found in animals, yeast, fungi and many bacteria where they play important basic biological functions in lipid metabolism and cell signaling and contribute to membrane fluidity in response to temperature fluctuation (Sperling et al. 2003). In the past decades, the integration of molecular and phylogenetic approaches has greatly advanced our understanding of the function of FAD genes in the biosynthesis of mono- and poly-unsaturated fatty acids in a range of organisms. Moreover, an ever-increasing number of acyl-CoA desaturase genes have been functionally characterized, demonstrating mechanistically their crucial role in the biosynthesis of pheromone and semiochemicals in *Drosophila* fruitflies, bees, wasps, beetles, and lepidopteran species. Variation in the number and expression of acyl-CoA desaturase genes have been shown to affect the diversity of pheromone signals between closely related species (Takahashi et al. 2001; Roelofs et al. 2002; Fang et al. 2009; Shirangi et al. 2009; Albre et al. 2012). The family is characterized by multiple episodes of expansion and contraction that occurred during the evolution of insects (Roelofs and Rooney 2003; Helmkampf et al. 2015). Consequently, the FAD gene family has become a model to study how structural and regulatory changes act in concert to produce new phenotypes.

Among the taxa available to study the molecular basis of pheromone production in an evolutionary framework leafroller moths (Lepidoptera: Tortricidae) provide a model system of choice (Roe et al. 2009). Leafrollers represent one of the largest families in the Lepidoptera with over 10,000 described species (Gilligan et al. 2018). Their larvae feed as leaf rollers, leaf webbers, leaf miners or borers in plant stems, roots, fruits or seeds. Many tortricid species are important pests and, due to their economic impact on human society, became the target of countless pheromone identification studies. In 1982, Roelofs and Brown published a comprehensive review on pheromones and evolutionary relationships among Tortricidae (Roelofs and Brown 1982). These authors related the patterns in pheromone diversity in a biosynthetic perspective to different postulated phylogenies of the Tortricidae, and specifically the two major group within Tortricidae, Tortricinae and Olethreutinae. Based on the pheromone identifications available at the time, it was suggested that species in the Tortricinae use mostly 14-carbon pheromone components (acetates, alcohols and aldehydes) whereas species in the Olethreutinae subfamily use mostly 12-carbon compounds. Building on recent advances towards a robust molecular phylogeny of Tortricidae (Regier et al. 2012; Fagua et al. 2017) and incorporating the information for the pheromones identified in 179 species and sex attractants reported for an additional 357 species we show that the proposed dichotomy is well-supported by the data currently available (Fig. 1). Furthermore, a majority of species in Tortricinae use pheromone components with double bonds in uneven positions, i.e., Δ9 isomers or Δ11 isomers. By contrast, the Olethreutinae pheromones typically contain components with double bonds in even positions (Δ8, Δ10), with doubly unsaturated Δ8Δ10:12C fatty acyl chains being one of the signature structures of the subfamily. Roelofs and Brown (1982) suggested that the use of a Lepidoptera-specific Δ11-desaturase acting on myristic acid (C_14_) could account biosynthetically for most of the pheromone compounds found in Tortricinae. This hypothesis was later confirmed with the functional characterization of FADs expressed in the female pheromone gland of several Tortricinae representatives. These include a desaturase that makes only the *E*11-isomer in the light brown apple moth *Epiphyas postvittana* (E11-14, E11-16 and E9E11-14)(Liu et al. 2002a), as well as desaturases from the redbanded leafroller moth *Agryrotaenia velutinana* (Liu et al. 2002b) and the obliquebanded leafroller moth *Choristoneura rosaceana* (Hao et al. 2002a) which both produce a mixture of Z/E11-14:Acids. In the case of Olethreutinae, Δ11-desaturation followed by chain-shortening could account for the Δ9:12C compounds found in several tribes of Olethreutinae. On the other hand, the biochemical pathways leading to the pheromone components with double bonds in even positions, the Δ8 and Δ10 as well as the doubly unsaturated Δ8Δ10:12C compounds in the Olethreutinae was not obvious.

**Figure 1:**
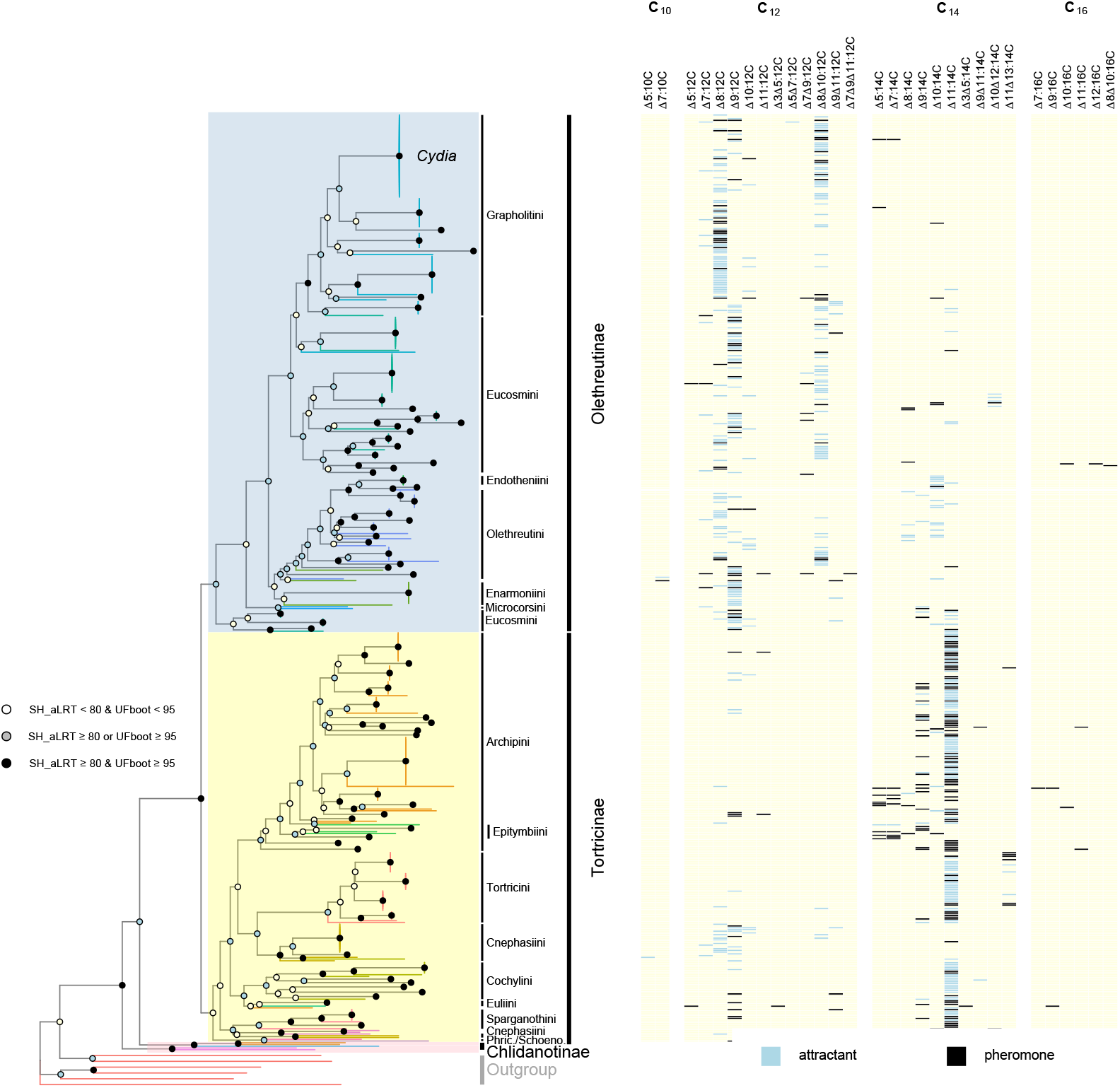
Phylogeny of Tortricidae and their associated female sex pheromone components. (left) The maximum likelihood tree was obtained for predicted nucleotide sequences of Tortricidae species (7591 aligned positions). The species represented comprise all tortricids for which pheromones or attractants have been reported plus some outgroups. Typically, one representative species was chosen per genus and contributed molecular evidence for all species in the genus. Outgroup species are represented by red branches whereas tortricid species from the same tribe are represented by branches of the same colour. Branch support values were calculated from 1000 replicates using the Shimodaira-Hasegawa-like approximate ratio test (SH_aLRT) and ultrafast bootstraping (UFboot). Support values for branches are indicated by colored circles, with color assigned based on thresholds of branch selection for SH-aLRT (80%) and UFBoot (95%) supports, respectively. The major subfamilies and represented tribe names are indicated (Phric: Phricanthini; Schoeno: Schoenotenini). (right) Heatmap indicating presence/absence of unsaturated fatty acid structure in bioactive molecules. Attractants correspond to compounds found to be attractive in either field or laboratory experiments; pheromone components correspond to sex attractants produced naturally by the organism and with a demonstrated biological activity on conspecific males. Double bond positions are annotated in Δ-nomenclature without referring to the geometry. Molecular and trait data retrieved from GenBank and the Pherobase, respectively.

The codling moth *Cydia pomonella* (Linnaeus) (Tortricidae: Olethreutinae: Grapholitini) is one of the most devastating pests in apple and pear orchards worldwide (Witzgall et al. 2008). With the goal of disrupting its reproduction, it has received substantial attention and been at the center of numerous studies focused on characterizing its communication system via sex pheromones. Its pheromone, (*E*8,*E*10)-dodecadien-1-ol, also known under the common name *codlemone*, was first identified using gas chromatography in combination with electroantennogram (EAG) recordings (Roelofs et al. 1971). This identification was later confirmed by fine chemical analysis (Beroza et al. 1974; McDonough and Moffitt 1974). The codling moth thus provides a relevant system to unravel the molecular pathway associated with the production of the Δ8Δ10:12C typical of Olethreutines. Previous studies support the hypothesis that the biosynthesis involves the desaturation of a Δ9 monoene intermediate. First, Arn et al. (1985) reported the presence of the unusual (*E*)-9-dodecenol (E9-12:OH) at about 10% of the doubly unsaturated alcohol in pheromone gland extracts and effluvia from *C. pomonella* females. Although this monoene does not carry any behavioral activity, its occurrence suggested that E9-12:Acyl could be an intermediate of codlemone biosynthesis in a process analogous to the biosynthesis of 10,12-dienic systems via Δ11 monoene intermediates observed in *Bombyx mori, Manduca sexta*, and *Spodoptera littoralis* (Yamaoka et al. 1984; Tumlinson et al. 1996; Navarro et al. 1997). The selective incorporation of deuterium-labeled E9-12 fatty acid precursors into codlemone and its direct precursor, *E*8,*E*10-dodecadienoate (E8E10-12:Acyl), supported the presence of an unusual Δ9 desaturase in *C. pomonella* and the biosynthesis of a conjugated diene system via 1,4-desaturation and the characteristic elimination of two hydrogen atoms at the allylic position of the double bond in the monoene intermediate precursor (Löfstedt and Bengtsson 1988)(Fig. 2). Similar conclusions were reached from a replicate study using *C. splendana* and *C. nigricana* females (Witzgall et al. 1996). To date, the FAD(s) central to the biosynthesis of codlemone and related fatty acyl derivates with a Δ8Δ10 system has not been characterized.

**Figure 2:**
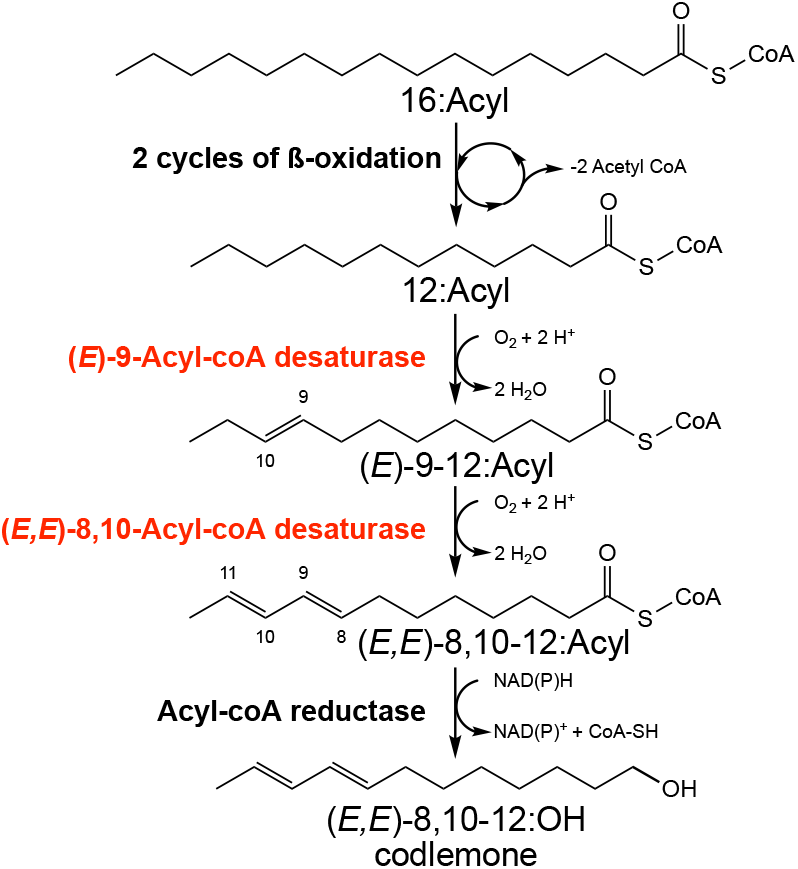
Pathway of codlemone biosynthesis. Palmitic acyl (C_16_) is first formed by the fatty acid synthesis pathway. Lauric acyl (C_12_) is then produced following two recurring reactions of beta-oxidation. A first desaturation occurs, introducing an *E*9 double bond between carbon 9 and 10 from the carboxyl terminus. The resulting monoenoic fatty acyl is then the substrate of a second desaturation via the removal of hydrogen atoms at the carbons 8 and 11, causing the formation of an *E*8*E1*0 conjugated diene system. Finally, an unknown fatty acyl-CoA reductase converts the fatty acyl precursor into the fatty alcohol (*E*8,*E*10)-dodecadienol commonly known as codlemone. The steps characterized in this study are highlighted in red.

The availability of a high-quality draft genome for *C. pomonella* provides an opportunity to comprehensively annotate and analyze relevant genes (Wan et al. 2019). Here we annotated a total of 27 FAD genes and performed a phylogenetic analysis, revealing expansions of different ages in the Δ9(KPSE) clade and in the Δ10/Δ11(XXXQ/E) clade. Using a transcriptomic approach, we determine the breadth of expression of all FAD genes and identified genes with upregulated expression in the pheromone gland of female *C. pomonella*, where the biosynthesis of codlemone takes place. We tested the function of these desaturases in heterologous expression systems and identified one FAD gene conferring on the cells the dual desaturase functions playing a key role in the biosynthesis of (*E8,E*10)-dodecadienol in *C. pomonella* and a signature structure of many Olethreutinae moth pheromones.

## Results

### Expansion of FADs in the Genome of C. pomonella

We identified candidates potentially involved in fatty acid synthesis by searching in the genome of *C. pomonella* for genes encoding fatty acid desaturases, which are characterized by a fatty-acid desaturase type 1 domain (PFAM domain PF00487). First, we improved the genome annotation by incorporating data from the pheromone gland transcriptome, a tissue which was not part of the panel of tissues used to generate the available annotation of *C. pomonella*. Searching our improved annotation, we identified 27 genes harboring the signature domain of FADs. While the vast majority of genes identified in our exhaustive search contain open-reading frames of 300 amino acids or longer, a small number of genes (i.e. 2-3) may represent pseudogenes or assembly errors. We adopted the nomenclature proposed by Knipple et al. (2002) in which genes are named based on the composition of 4 amino acid residues at a signature motif. Next, we looked at their genomic organization. We found that FAD genes are distributed across 12 of the 27 autosomes, with no FAD genes on either Z or W sex chromosome (chr1 and chr29, respectively) (Fig. 3). Two genes, Cpo_QPVE and Cpo_MATD(2), were found on unplaced scaffolds. As is typical for members of a multigene family, we identify several clusters corresponding to tandemly duplicated genes. Moreover, several genes are present as the single member of the family on a given autosome.

**Figure 3:**
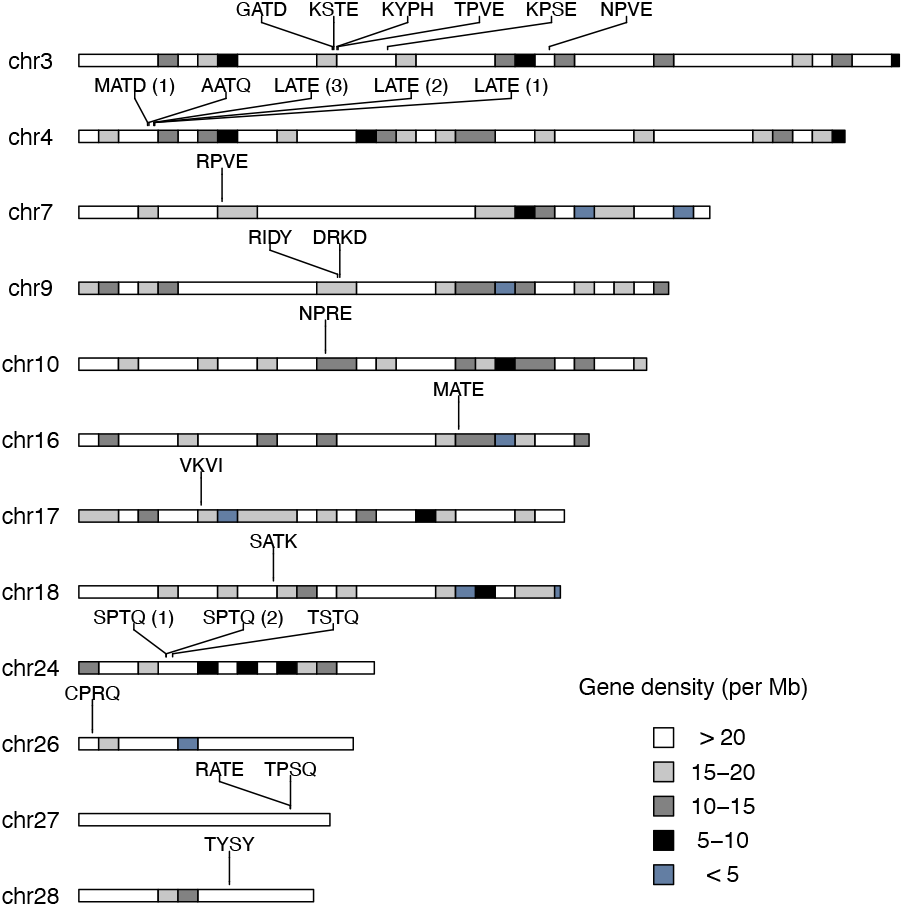
Genomic organization of FAD genes in *Cydia pomonella*. The position of the 27 predicted FAD genes were mapped to the genome. 25 genes could be placed on 12 autosomes; 2 genes, Cpo_MATD(2) and Cpo_QPVE, are located on unplaced scaffolds (not drawn). The distribution analysis showed that there are 5 gene clusters which contain 2 or more FAD genes. Names refer to chromosome names in the genome assembly. Chromosomes are drawn to scale and the bands represent gene density.

### Phylogenetic Analyses Place Expansions in the Δ9(KPSE) and Δ10/Δ11(XXXQ/E) Clades

In order to determine which of the genes identified above could be important for codlemone biosynthesis, we assessed the functional classes represented by these genes using phylogenetic analyses. To that end, we aligned the predicted protein sequences of the *C. pomonella* genes with a panel of FAD sequences from Lepidoptera species including genes for which function have been previously characterized using *in vitro* heterologous expression. We found representatives of the eight insect acyl-CoA desaturase subfamilies *sensu* Helmkampf et al. (Helmkampf et al. 2015) (Fig.4, Fig. S1).

**Figure 4:**
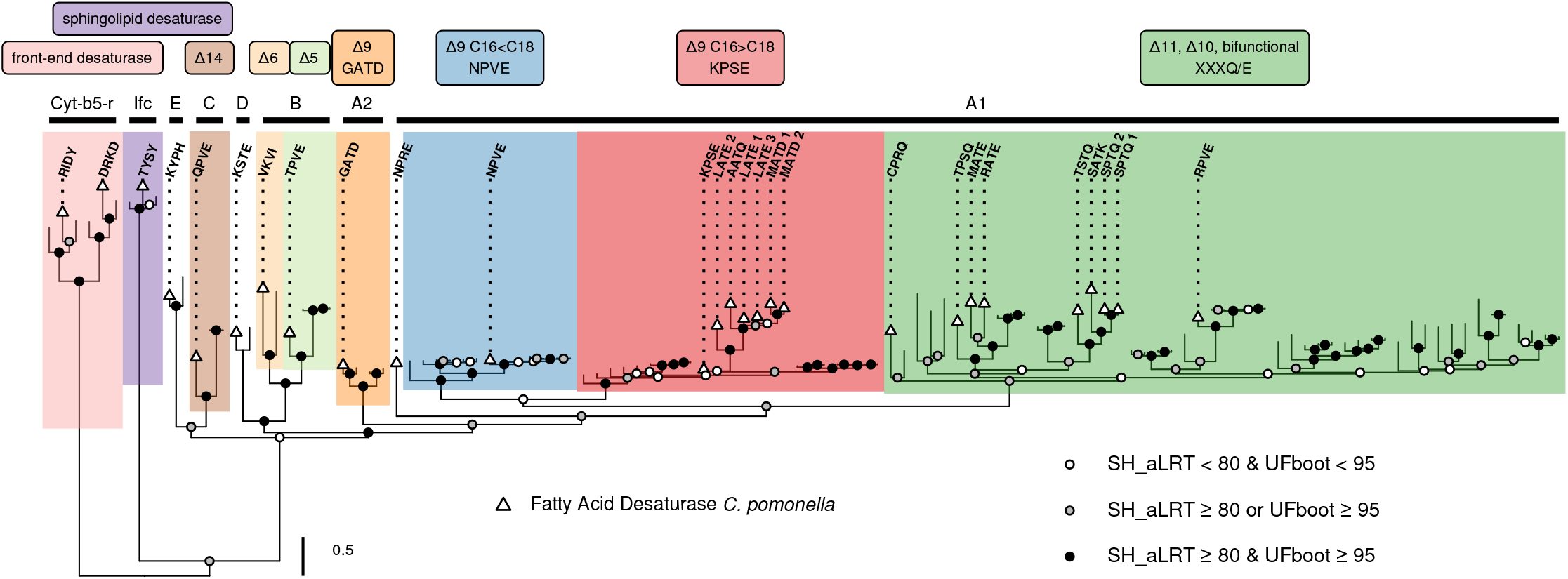
Phylogeny of lepidoptera FAD genes. The maximum likelihood tree was obtained for predicted amino acid sequence of 114 FAD genes (805 aligned positions) of 28 species, with branch support values calculated from 1000 replicates using the Shimodaira-Hasegawa-like approximate ratio test (SH_aLRT) and ultrafast bootstraping (UFboot). Support values for branches are indicated by filled circles, with the color assigned based on SH-aLRT and UFBoot supports using 80% and 95% as thresholds of branch selection for SH-aLRT and UFBoot supports, respectively. The major constituent six subfamilies of First Desaturase (A1 to E) and two subfamilies of Front-End (Cyt-b5-r) and Sphingolipid Desaturases (Ifc), respectively, are indicated following the nomenclature proposed by Helmkampf et al. (2015). For First Desaturases, the different shades correspond to the indicated putative biochemical activities and consensus signature motif (if any). Triangles indicate sequences from *C. pomonella* (see also Fig. S1 for an extended version of this tree). The scale bar represents 0.5 substitutions per amino acid position.

All subfamilies form highly supported groups, with the exception of the relationship between Desat A1 and A2, which appear weakly or strongly supported, depending on the set of genes used in our analyses. With a characteristic domain architecture (PF08557 in front of PF00487), Cpo_TYSY encodes a putative Sphingolipid Delta-4 desaturase and is homologous to *interfertile crescent* in *D. melanogaster* (Ifc). Two tandemly duplicated genes, Cpo_RIDY and Cpo_DRKD, contain a Cytochrome b5-like heme binding domain (PF00173) in front of the fatty acid desaturase domain and group with putative Front-End desaturases homologous to *Cytochrome b5-related* in *D. melanogaster* (Cyt-b5-r). These three genes bear little similarity with the other 24 FAD genes which encode First Desaturases of the subfamilies Desat A1 through E.

Desat C, D, and E are present as single-copy genes in *C. pomonella* (Cpo_QPVE, Cpo_KSTE, and Cpo_KYPH, respectively). Cpo_QPVE groups with the Δ14 desaturase identified in male and female corn borer pheromone biosynthesis (Lepidoptera: Crambidae) (Roelofs et al. 2002; Lassance and Löfstedt 2009).

Desat B, which is particularly expanded in Hymenopterans and in *Bombyx mori* (Helmkampf et al. 2015), has only two representatives in *C. pomonella*. These genes, Cpo_VKVI and Cpo_TPVE, are homologous to the *E*6 desaturase identified in *Antheraea pernyi* (Lepidoptera: Saturnidae) (Wang et al. 2010) and the *Z*5 desaturase identified in *Ctenopseustis obliquana* (Lepidoptera: Tortricidae) (Hagström et al. 2014), respectively.

Desat A2 subfamily is characterized by a single-copy gene, Cpo_GATD, which seem to be typical of Lepidoptera. The homolog identified in *Choristoneura parallela* (Lepidoptera: Tortricidae) is associated with the formation of Δ9 desaturation in saturated acyl moieties of a range of length (14C-26C) (Liu et al. 2004).

Finally, Desat A1 forms the largest group and experienced a particularly dynamic evolutionary history. With 17 genes *C. pomonella* harbors twice as many genes as *Bombyx mori* (7 in the most recent version of the genome SilkDB 3.0 (Lu et al. 2020)). While Δ9 C16<C18 (NPVE) is represented by a single-copy gene, we found two significant expansions in the group corresponding to Δ9 C16>C18 (KPSE) and Δ11, Δ10 & bifunctional (XXXQ/E) FADs. The tandemly arrayed cluster found in chr4 correspond to an extension within the A1 subfamily encoding Δ9 C16>C18 FADs. Nine genes are found in the Lepidoptera-specific A1 subfamily encoding Δ11, Δ10 & bifunctional FADs. These genes are scattered across 6 private chromosomes, none of which containing members of the other functional classes.

Based on their placement in the FAD gene tree, these genes can be sub-divided into four groups (Fig.4, Fig. S1). Group1 contains Cpo_SPTQ(1), Cpo_SPTQ(2), Cpo_SATK and Cpo_TSTQ, which are homologous to the Δ10 desaturase identified in the New Zealand tortricid *Planotortrix octo* (Lepidoptera: Tortricidae) (Hao et al. 2002). Group2 is formed with Cpo_RATE, Cpo_MATE, and Cpo_TPSQ, which are homologous to the FAD with Δ6 activity isolated from *Ctenopseutis herana* (Lepidoptera: Tortricidae) (Albre et al. 2012). Group3 is represented by Cpo_RPVE, which groups with desat2 from *Ctenopseutis obliquana* and allied species (Lepidoptera: Tortricidae) (Albre et al. 2012) and Lca_KPVQ from *Lampronia capitella* (Lepidoptera: Prodoxidae) (Liénard et al. 2008). Finally, Cpo_CPRQ falls near Dpu_LPAE from *Dendrolimus punctatus* (Lepidoptera: Lasiocampidae) (Liénard et al. 2010).Based on our phylogenetic reconstructions, several genes are plausible candidates in the biosynthesis of codlemone. First, genes from the Δ9 C16>C18 (KPSE) subfamily have been implicated in pheromone biosynthesis and carry activities similar to those expected in *C. pomonella*. Specifically, the Dpu_KPSE from *Dendrolimus punctatus* (Lepidoptera: Lasiocampidae) produces a range of Δ9-monounsaturated products including Z9- and E9-12:Acyl (Liénard et al. 2010). This exemplifies that enzymes in this subfamily can introduce *cis* and *trans* double bonds in lauric acid (12C). Furthermore, when supplemented with Z7- and E7-14:Acyl, Dpu_KPSE can introduce a second desaturation to produce Δ7Δ9-14:Acyl, illustrating the formation of conjugated double bonds by this type of FAD. The expansion of the KPSE clade we report in *C. pomonella* is an interesting coincidence. Second, the abundance of representatives in the Δ11, Δ10 & bifunctional (XXXQ/E) subfamily support the possibility that one or several could be involved in codlemone biosynthesis. This group contains many genes involved in the biosynthesis of conjugated double bonds, e.g. Bmo_KATQ in *Bombyx mori* (Lepidoptera: Bombycidae) (Moto et al. 2004), Mse_APTQ in *Manduca sexta* (Lepidoptera: Sphingidae) (Matoušková et al. 2007), Lca_KPVQ in *Lampronia capitella* (Lepidoptera: Prodoxidae) (Liénard et al. 2008), and Dpu_LPAE in *Dendrolimus punctatus* (Liénard et al. 2010). Cpo_RPVE groups with Lca_KPVQ, which encodes an enzyme involved in the desaturation of 16:Acyl and Z9-14:Acyl to produce the conjugated Z9Z11-14:Acyl pheromone precursor of *L. capitella* (Liénard et al. 2008). Another interesting candidate is Cpo_CPRQ, which clusters near Dpu_LPAE, an enzyme capable of catalyzing the production of E9Z11-16:Acyl and E9E11-16:Acyl (Liénard et al. 2010).

### Patterns of Gene Expression Identify Cpo_CPRQ and Cpo_SPTQ(1) as Candidates

To identify candidates for codlemone production we compared the expression levels of the twenty-seven genes found in the genome using published RNA sequencing (RNA-Seq) data from various tissues and life-stages augmented with two new pheromone gland data sets (this study). Since codlemone is found in the pheromone gland of females, which is located at the tip of the abdomen, we hypothesized that its desaturase(s) would be enriched or even exclusively expressed in this sex and in the population of cells forming this tissue.

We found strong qualitative differences in the transcription profile of the 27 examined FAD genes (Fig. 5). The genes divide into three clusters. The first two correspond to genes that appear ubiquitously expressed. These include the two putative Front-End desaturases Cpo_DRKD and Cpo_RIDY (Cyt-b5-r), the Sphingolipid Δ4 desaturase Cpo_TYSY (Ifc), and five First desaturases, namely Cpo_NPVE (A1; Δ9 C16<C18), Cpo_KPSE (A1; Δ9 C16>C18), Cpo_RPVE (A1; Δ11/Δ10), Cpo_KSTE (D), Cpo_KYPH (E). With the exception of Cpo_RPVE, these genes exhibit a high evolutionary stability characterized by a single-copy gene with rare cases of gains and losses across insects. Their broad expression profiles suggest that these genes fulfill a fundamental metabolic function, making them unlikely candidates for pheromone biosynthesis. By contrast, the other First Desaturase genes found in the third cluster are characterized by a sparse expression profile and greater tissue and/or life-stage specificity: head: Cpo_TPSQ (A1; Δ11/Δ10); ovary/female abdomen: Cpo_TPVE (B; Δ5); pheromone gland: Cpo_CPRQ (A1; Δ11/Δ10), Cpo_SPTQ(1) (A1; Δ11/Δ10); eggs: Cpo_NPRE (A1), Cpo_VKVI (B; Δ6). These expression patterns are suggestive of more specialized functions. Specifically, their expression profiles and levels indicated that Cpo_CPRQ and Cpo_SPTQ(1) (average FPKM in pheromone gland: 6082.0 and 1451.5, respectively) represent primary candidates for functional testing.

**Figure 5:**
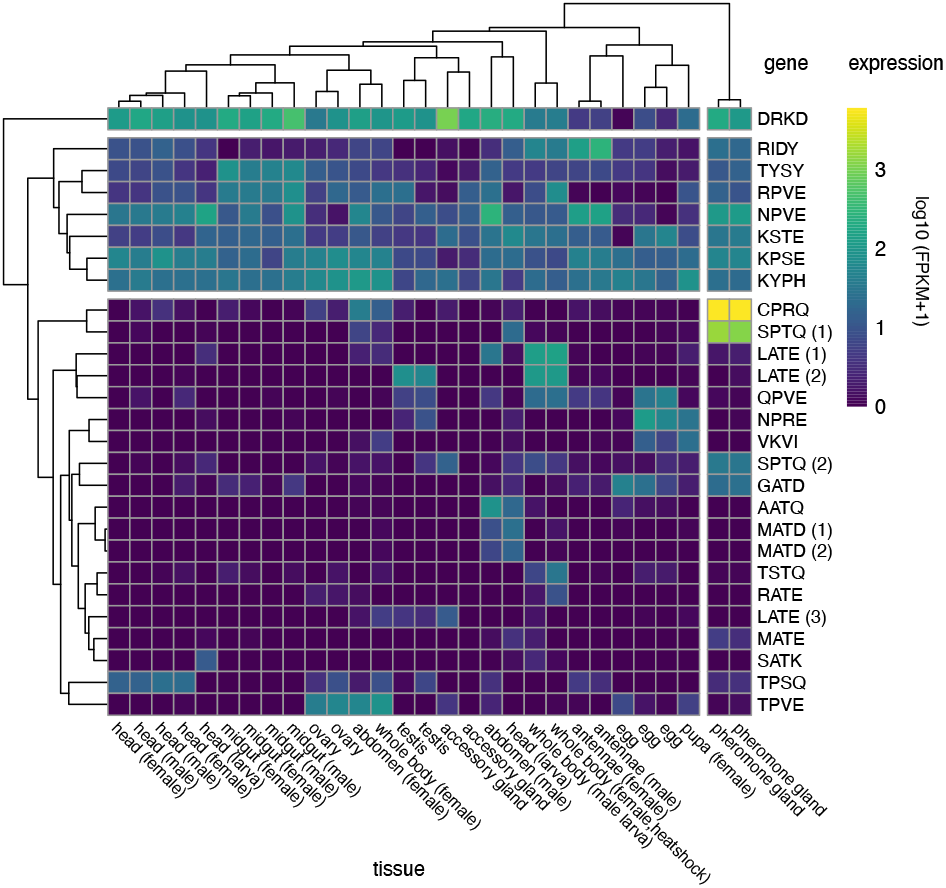
Expression profiling of FAD genes in different tissues of *Cydia pomonella*. The heatmap represents the absolute expression value (log FPKM) of all FAD genes in the corresponding tissues. Genes are identified by their signature motif. Automatic hierarchical clustering of FAD genes distinguishes three clusters: cluster I and II contain genes with ubiquitous expression across tissues and developmental stages; cluster III contain genes with more divergent expression and higher tissue specificity. Automatic hierarchical clustering of tissues indicates that pheromone gland samples have a distinguishable profile, which is characterized by the over-expression of Cpo_CPRQ and Cpo_SPTQ(1). Estimates of abundance values were obtained by mapping reads against the genome. The NCBI SRA accession numbers of all RNA-Seq data sets used are given in Supplementary Table S4.

### Functional Characterization Demonstrates the E9 FAD Activity of Cpo_CPRQ and Cpo_SPTQ(1)

Previous studies have shown that the baker’s yeast *Saccharomyces cerevisiae* is a convenient system for studying *in vivo* the function of FAD genes and it has become the eukaryotic host of choice for the heterologous expression of desaturases from diverse sources including moths. Conveniently, yeast cells readily incorporate exogenous fatty acid methyl esters and convert them to the appropriate coenzyme A thioester substrates required by FADs.

First, we cloned Cpo_CPRQ and Cpo_SPTQ(1) into a copper-inducible yeast expression vector to generate heterologous expression in *S. cerevisiae*. We carried out assays with precursor supplementation to ensure availably of the medium-chain saturated fatty acids, i.e. lauric (C_12_) and myristic acids (C_14_), which are typically less abundant. Cpo_CPRQ conferred on the yeast the ability to produce a small amount of E9-12:Acyl and had no detectable activity outside of the C_12_ substrate (Fig. 6). Cpo_SPTQ(1) showed Δ9 desaturase activity, producing E9- and Z9-12:Acyl, Z9-14:Acyl and Z9-16:Acyl. Double bond positions in these products were confirmed by dimethyl disulfide (DMDS) derivatization (Fig. 6). In addition to retention times matching those of synthetic standards, spectra of the DMDS derivatives of the methyl esters displayed the following diagnostic ions: Z9-16:Me (M^+^, m/z 362; A^+^, m/z 145; B^+^, m/z 217), Z9-14:Me (M^+^, m/z 334; A^+^, m/z 117; B^+^, m/z 217), E9- and Z9-12:Me (M^+^, m/z 306; A^+^, m/z 89; B^+^, m/z 217). No doubly unsaturated products were detected in these experiments.

**Figure 6:**
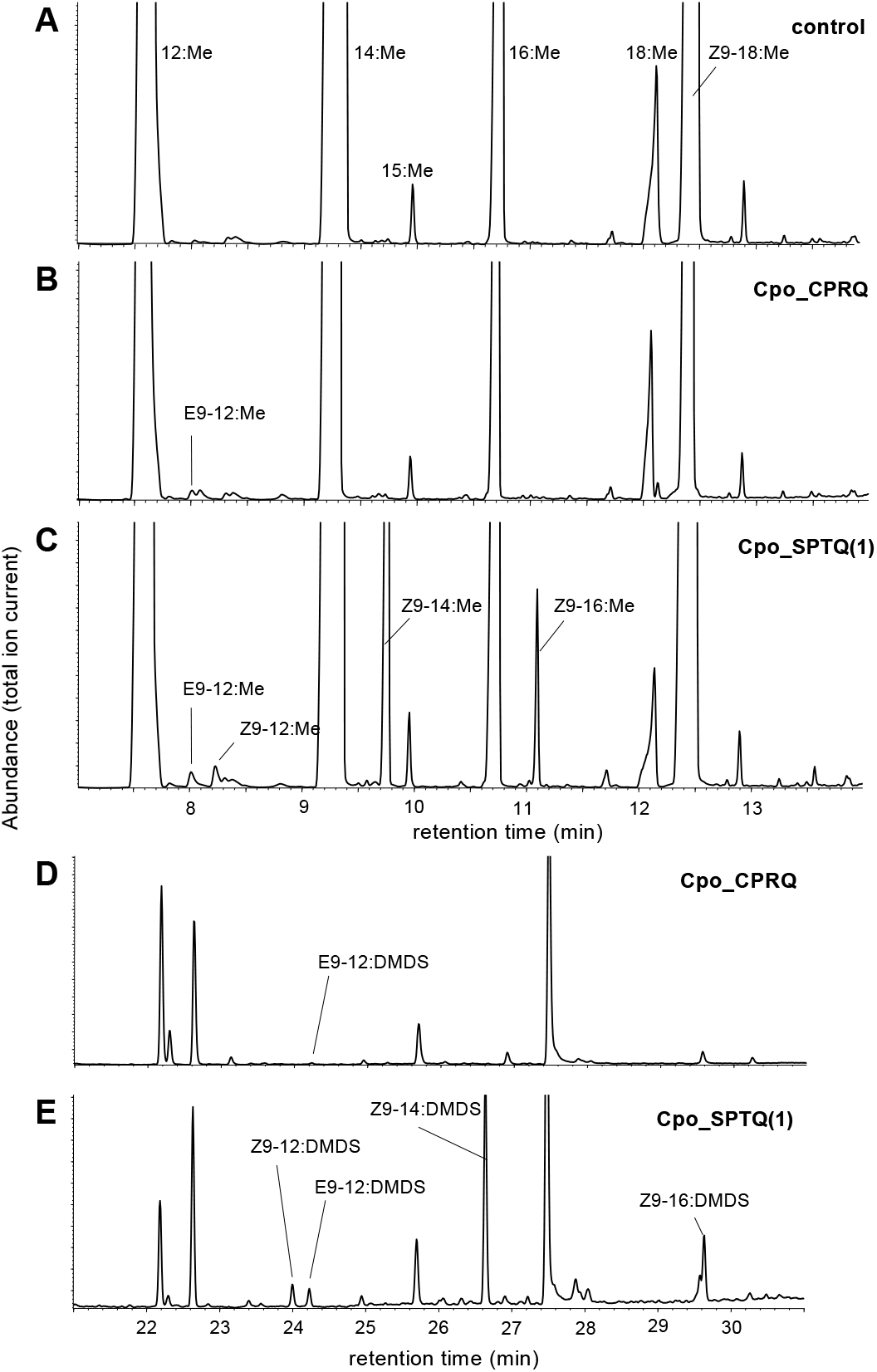
Functional characterization of desaturase activity of candidate genes in yeast. Total ion chromatograms of fatty acid methyl ester (FAME) products of Cu2+-induced ole1 elo1 S. cerevisiae yeast supplemented with saturated acyl precursors and transformed with (A) empty expression vector (control), (B) pYEX-CHT-Cpo_CPRQ and (C) pYEX-CHT-Cpo_SPTQ(1). Confirmation of identity of enzyme products was obtained by comparison of retention time, mass spectra of synthetic standards. Confirmation of double bond positions by mass spectra of DMDS adducts (D, E).

In a second round of experiments we assayed 7 other First desaturases that could be amplified from pheromone gland cDNA. As predicted from their respective placement in the FAD phylogeny, Cpo_NPVE, Cpo_KPSE and Cpo_GATD encode Δ9 desaturases (Fig. S2). Cpo_NPVE is a Δ9 desaturase with preference for myristic (C_14_) and palmitic (C_16_) acid, showing only weak activity on lauric acid (C_12_). Cpo_GATD and Cpo_KPSE showed Δ9 desaturase activities mainly on C_16_ but also some activity on C_14_. The other 4 desaturases had no detectable activity in the yeast system.

Finally, we supplemented the growth medium with the monounsaturated methyl esters E9-12:Me and Z9-12:Me. The doubly unsaturated codlemone precursor, E8E10-12:Acyl, was inconsistently detected in some replicates of the assays with Cpo_CPRQ. No doubly unsaturated products were detected with the other constructs.

### Functional Analysis Identify Cpo_CPRQ as the FAD Catalyzing Δ8Δ10 Desaturation

In order to evaluate the presumptive Δ8Δ10 desaturase activity of Cpo_CPRQ, we further investigated the activity of the enzyme in an insect cell line. We reasoned that the inconsistent activity of the enzyme in yeast could be due to the cellular environment of the expression system. When we expressed Cpo_CPRQ using the Sf9 cell system and supplementing the culture medium with lauric acid, we could observe the consistent production of E9-12 as well as E8E10-12, demonstrating that Cpo_CPRQ catalyzes the biosynthesis of E8E10-12:Acyl and its monounsaturated intermediate E9-12:Acyl (Fig. 7). As expected from the previous experiments in yeast, we did not detect activity on longer chain substrates. The retention time and mass spectrum of the E8E10-12:Me peak were identical to those of the synthetic standard. Double bond positions of the conjugated system were confirmed by derivatization with 4-methyl-1,2,4-triazoline-3,5-dione (MTAD)(Fig. 7). Finally, we carried out assays to further evaluate the stereospecificity of the enzyme. When provided with E9-12:Me, Cpo_CPRQ produced the doubly unsaturated product (Fig 7). In contrast, we found no evidence for activity on the corresponding diastereomer, Z9-12:Me (Fig 7).

**Figure 7:**
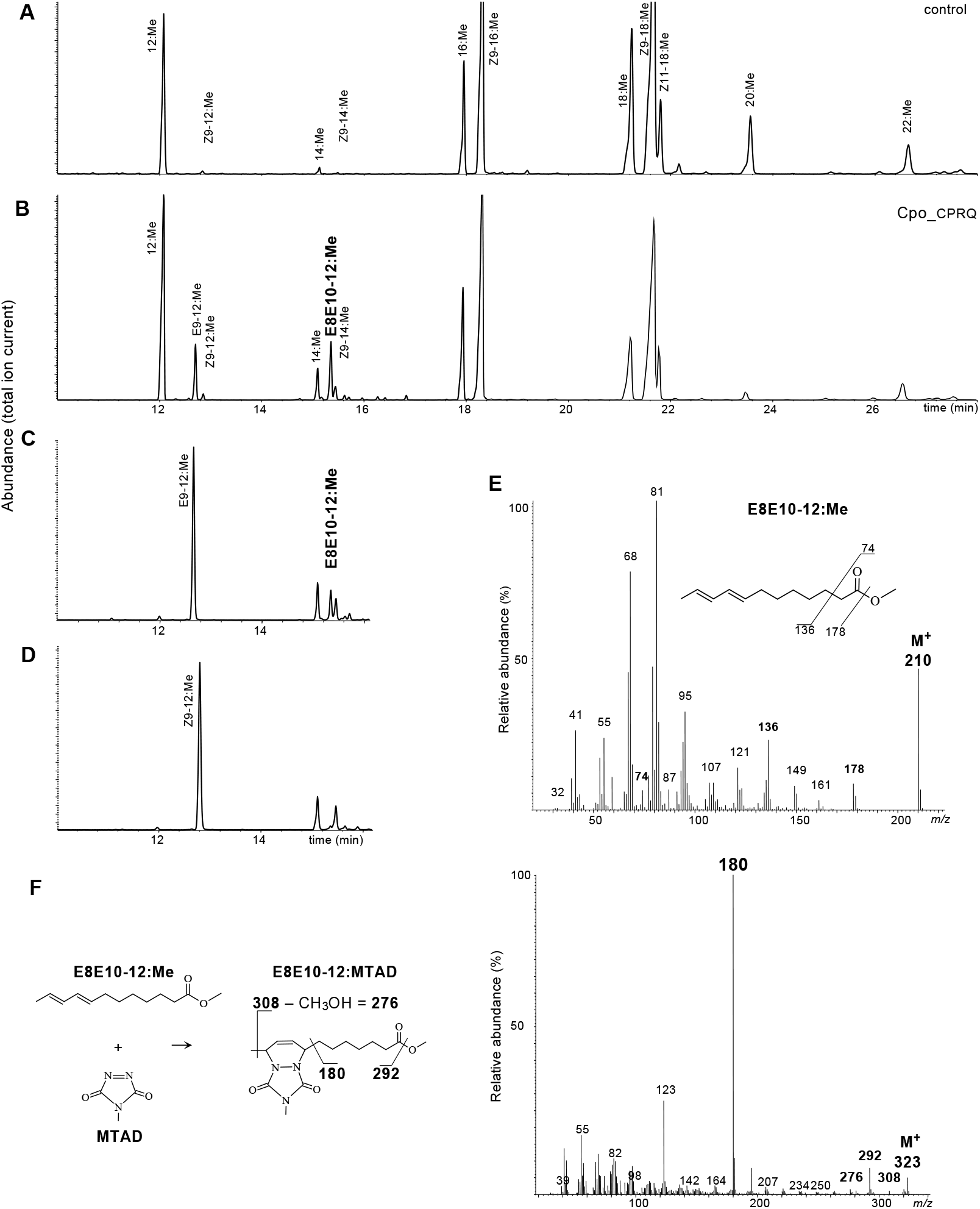
Functional characterization of desaturase activity of Cpo_CPRQ in insect cells. Total ion chromatograms of methyl esters (FAME) samples from Sf9 cells supplemented with lauric methyl ester (C12) infected with (A) empty virus (control) or (B) recombinant baculovirus expressing Cpo_CPRQ. Cpo_CPRQ produces large amount of E9-12:Me and E8E10-12:Me. Sf9 insect cells infected with bacmid expressing Cpo_CPRQ in medium in the presence of the monoenic intermediate (C) (*E*)-9-dodecenoic methyl ester (E9-12:Me) or (D) (*Z*)-9-dodecenoic methyl ester (Z9-12:Me). The retention time and mass spectrum of the E8E10-12:Me peaks observed after addition of 12:Me and E9-12:Me were identical with those of the synthetic standard. The relatively long retention time (compound eluting later than 14:Me) is in agreement with what is expected from a diene with conjugated double bonds. (E) Spectrum of the E8E10-12:Me peak in panel B. The relatively abundant molecular ion m/z 210 is in agreement with the expectation for a diene with conjugated double bonds. (F) Analyses of MTAD-derivatized samples displayed diagnostic ions at m/z 323 (M+), m/z 308 and m/z 180 (base peak), confirming the identification of the conjugated double bonds system.

Altogether, these results show that Cpo_CPRQ exhibit both the ability to catalyze the *trans* desaturation of lauric acid to form E9-12:Acyl and the 1,4-desaturation of the latter to form the E8E10-12 conjugated pheromone precursor of codlemone.

## Discussion

In this study, we show the key role of a desaturase with a dual function in the biosynthesis of codlemone. We confirm the results suggested by the labelling experiments reported by Löfstedt and Bengtsson (1988) and demonstrate that Cpo_CPRQ can conduct both the initial desaturation of 12:Acyl into E9-12:Acyl, and then transform E9-12:Acyl into E8E10-12:Acyl, the immediate fatty acyl precursor of E8E10-12:OH, the sex pheromone of *C. pomonella* (Fig. 2). Phylogenetically, Cpo_CPRQ clusters in the Lepidoptera-specific XXXQ/E clade (Fig 4). Several FADs found in that clade have been demonstrated to be involved in the pheromone biosynthesis of many moth species, including the species of Tortricinae using Δ11 (*Epiphyas postvittana, Choristoneura rosaceana, Agryrotaenia velutinana*) and Δ10 desaturations (*Planotortrix octo*) (Liu et al 2002a; Hao et al. 2002a; Liu et al 2002b; Hao et al. 2002b). Our phylogenetic reconstruction indicates that Cpo_CPRQ does not appear orthologous to any of those genes (Fig. S1), suggesting the recruitment of a different ancestral duplicate early in the evolution of Olethreutinae. In addition to Cpo_CPRQ, Cpo_SPTQ(1) clustered in the same clade and had a similar activity profile on lauric acid, although it produces more Z9-than E9-12:Acyl. With an expression at about 25% the level of Cpo_CPRQ, Cpo_SPTQ(1) is the second most highly expressed FAD gene in the pheromone gland, a tissue in which this FAD appear exclusively expressed. Although Cpo_CPRQ appears sufficient to produce E8E10-12:Acyl, we cannot exclude the possibility that Cpo_SPTQ(1) contributes to the production of the monoene intermediate and is involved in the pheromone biosynthesis of codlemone or E9-12:OH, a pheromone component eliciting antennal response but with no apparent behavioral effect (Witzgall et al. 2001). Interestingly, even though they are phylogenetically clustered within the Δ11/Δ10-desaturase subfamily, Cpo_CPRQ and Cpo_SPTQ(1) exhibit primarily Δ9-desaturase activity, which likely represent the ancestral state. Whether this represents a conservation or a reversal to the ancestral function awaits further investigations. This finding provides exciting opportunities to study the structure-function relationships in FADs, in particular the structural determinant of regioselectivity.

Expansion of gene families is usually regarded as playing a key role in contributing to phenotypic diversity. The acyl-CoA desaturase gene family is characterized by a highly-dynamic evolutionary history in insects (Roelofs and Rooney 2003; Helmkampf et al. 2015). Bursts of gene duplication have provided many opportunities for evolution to explore protein space and generate proteins with unusual and novel functions. In moths, many examples highlight the role played by the diversification in FAD functions in allowing the expansion of the multi-dimensional chemical space available for communication via sex pheromones. Careful re-annotation of the *C. pomonella* genome and phylogenetic analyses of identified desaturase genes allowed us to identify 27 genes representative of all desaturase subfamilies previously identified in insects. Interestingly, all functional classes previously identified in moths (i.e. Δ5, Δ6, Δ9, Δ10, Δ11, Δ14) have homologs in *C. pomonella*. This indicates that FAD genes tend to be preserved in the genomes for long periods of time. The availability of several chromosome-level assemblies in Lepidoptera has revealed that the genome architecture is generally conserved, even among distant species, with few to no structural rearrangements, i.e. inversions and translocations, being observed (but see (Hill et al. 2019)). This is attested by the high level of synteny existing between *C. pomonella* and *Spodoptera litura* (Lepidoptera: Noctuidae), two representatives of lineages which shared their last common ancestor ca. 120 MYA (Kawahara et al. 2019; Wan et al. 2019). This stability over long evolutionary time means that we can draw tentative conclusions from the genomic organization of gene families. Specifically, the genomic location of FAD genes in *C. pomonella* suggests that the diversification of this gene family is ancient. First Desaturase genes from all groups *sensu* Helmkampf et al. (2015) have representatives in species that have shared their last common ancestor over 300 MYA. Moreover, we found several expansions with differing patterns within Δ9 C16>C18 (KPSE) and the Lepidoptera-specific gene subfamily Δ11, Δ10 & bifunctional (XXXQ/E). Genes clustered in tandem arrays represent evolutionary recent duplications, as exemplified by the expansion in Δ9 C16>C18 (KPSE). These duplicates are likely the product of imperfect homologous recombination. By contrast, with 9 representatives spread across 6 chromosomes, the Δ11/Δ10 subfamily comprises both recent duplicates (e.g., Cpo_SPTQ(1) and Cpo_SPTQ(2)) as well as unlinked FAD genes (such as Cpo_CPRQ). The latter genes are likely to be the products of more ancient events of duplications and chromosomal rearrangements. The location of the majority of genes within this clade suggests that the diversification within this subfamily essential for the pheromone biosynthesis occurred early in the evolution of the Lepidoptera. As genome data from more species become available, it will be possible to follow the evolutionary trajectories of this gene family which has been a key contributor to pheromone evolution.

Expansion of gene families is often followed by a relaxation of selection, which translates into an acceleration of the mutation rate leading to the accumulation of non-synonymous nucleotide substitutions (Bielawski and Yang 2003; Taylor and Raes 2004). Looking at the impact of selection on desaturase genes in insects, Helmkapmf and co-authors (2015) found little evidence for positive selection but rather purifying selection acting on all desaturase genes in insects, even in expanded subfamilies. This could be indicative that large constraints along the coding sequences to not change sites affecting the proper enzymatic function outweigh the potential for diversifying selection. Modification of the regulation of duplicated genes is also expected, with some paralogs specializing to a limited set of tissues or developmental stages (Taylor and Raes 2004). Data from a panel representing a relatively large variety of tissue types and developmental stages reveal a large expression divergence between FAD genes. Interestingly, we see that genes present in recent expansion (i.e. Δ9 C16>C18 (KPSE)), of which members form tandem arrays in the genome, have low to no expression in the large panel of samples we analyzed. The same is also true for *Bombyx mori*. The evidence at hand suggest that these may have limited physiological relevance. This advocates for caution when interpreting the importance and functional consequences of expansions. Some expansions can contribute to lineage-specific adaptation and an increased demand for variability in associated traits. However, it is almost certain that many expansions will not reflect a response to changes and be responsible for the evolution of novel traits but rather have their evolutionary origin in the stochastic nature of the gene duplication process. Functional testing is paramount to advance our mechanistic understanding of the proximate consequences of variation in the size of multigene families.

Previously characterized FADs that catalyze the formation of conjugated fatty acids among moth sex pheromones precursors fall into two groups: (1) double bonds can be inserted via sequential 1,2-desaturation to produce a diene, either by the action of two different desaturases operating consecutively as postulated in *Dendrolimus punctatus* (Lasiocampidae) (Liénard et al., 2010), or by the same desaturase operating twice with an intermediate chain-shortening step as suggested for *Epiphyas postvittana* (Tortricidae) (Liu et al. 2002a), *L. capitella* (Liénard et al. 2008), and *Spodoptera littoralis* (Noctuidae) (Xia et al. 2019); (2) bifunctional desaturases can introduce the first double bond and then rearrange the monoene to produce a diene with conjugated double bonds via 1,4-desaturation, as seen in *Bombyx mori* (Bombycoidae) (Moto et al. 2004), *S. littoralis* (Noctuidae) (Serra et al. 2006) and *Manduca sexta* (Sphingidae) (Matoušková et al. 2007). The *C. pomonella* desaturase falls into the latter category: Cpo_CPRQ introduces first an *E*9 double bond in the 12:Acyl saturated substrate and then transforms the *E*9 unsaturation into the *E*8*E*10 conjugated double bonds. The independent evolution of similar features in enzymes used by distantly related species offers the opportunity to further study mechanistically the determinants of the switch between 1,2 and 1,4-dehydrogenation.

Recent molecular phylogenetic studies (Regier et al. 2012; Fagua et al. 2017) provide strong support for the monophyly of Tortricinae and Olethreutinae, the two most speciose subfamilies within Tortricidae, and the paraphyly of Chlidanotinae. Our tribal-level tree for Tortricidae combined with data on moth pheromones and attractants corroborate the pattern of pheromone components in Tortricinae and Olethreutinae reviewed and discussed by Roelofs and Brown (1982). Tortricinae species use mostly 14-carbon pheromone components with unsaturation introduced via Δ11 desaturation. Molecular and biochemical studies have confirmed the paramount role of Δ11 FADs in the pheromone biosynthesis of not only tortricines, but the vast majority of ditrysian Lepidoptera (see for instance (Jurenka 2003; Roelofs and Rooney 2003; Liénard et al. 2008, 2014; Liénard and Löfstedt 2010)). No pheromones are reported for the Chlidanotinae. However, the sex attractants reported for three species include Z9-14:OAc and Z11-16:OAc, which supports the idea that Δ11 desaturation in combination with chain-shortening may be typical of this subfamily. By contrast, the Olethreutinae subfamily use mainly 12-carbon compounds. The Δ9-12 chain structures used by many species could be accounted for the action of a Δ11 enzyme producing Δ11-14:Acyl as an intermediate following by chain-shortening to Δ9-12:Acyl. For example, the host races of the larch budworm *Zeiraphera diniana*(Tortricidae: Olethreutinae: Eucosmini) developing on larch (*Larix decidua*) or Cembra pine (*Pinus cembra*) used respectively E11-14:OAc and E9-12:OAc as pheromone (Guerin et al. 1984), and it is parsimonious to hypothesize that polymorphism in the activity of the chain-shortening enzyme is responsible for this difference between host races that show little to moderate genomic differentiation (Emelianov et al. 2004). Our data suggest however that an alternative pathway may exist. Indeed, two of the enzymes characterized in the present study (i.e. Cpo_SPTQ(1) and Cpo_CPRQ) can catalyze the production of Δ9-12:Acyl directly from lauric acid (C_12_), bypassing the need for prior Δ11 desaturation and chain-shortening. In addition to Δ9-12 chain structures, 12-carbon components with Δ8 and Δ8Δ10 systems are frequently used by olethreutines. Specifically, these occur in the three major tribes: Grapholitini, Eucosmini, and Olethreutini (Fig. 1). The identification of the bifunctional desaturase catalyzing the biosynthesis of Δ8Δ10-12 fatty acid derivatives sheds new light on the pathway associated with the production of pheromone of a large number of tortricids. Monounsaturated compounds with double bonds in Δ8 position frequently co-occur with conjugated Δ8Δ10, including in trace amount in *C. pomonella* (Witzgall et al. 2001). However, since we did not detect any Δ8 activity towards the lauric acid from Cpo_CPRQ or Cpo_SPTQ(1), we anticipate that a different FAD plays a role in the pheromone biosynthesis of Δ8 monoenes. This is supported by the pheromone bouquets identified for several species of olethreutines. In the podborer *Matsumuraeses falcana* (Tortricidae: Olethreutinae: Grapholitini), the pheromone consists of a mixture of E8-12:OAc and E8E10-12:OAc, plus minor amounts of (*E*7,*Z*9)- and (*E*7,*E*9)-dodecadienyl isomers, and the monoenes E10-12:OAc and E10-14:OAc (Wakamura 1985). The major pheromone component in *Hedya nubiferana* (Tortricidae: Olethreutinae: Olethreutini) is E8E10-12:OAc but in addition the pheromone contains relatively large amounts of Z8- and E8-12:OAc (Frerot et al. 1979). The same holds for *Epiblema foenella* (Tortricidae: Olethreutinae: Eucosmini) in which the major pheromone component was identified as Z8Z10-12:OAc: in addition to the other three geometric isomers of the diene, the females produce also E8- and Z8-12:OAc but no E9-12:OAc (Witzgall et al. 1996). Biosynthesis of these bouquets suggests the involvement of a FAD with Δ8 or Δ10 activity. A Δ10 FAD is the key enzyme involved in the production of monounsaturated compounds with double bonds in Δ8 position. Specifically, in *Planotrotrix sp*. and *Ctenopseustis sp*. (Tortricidae: Tortricinae: Archipini), Z8-14:OAc is bionsynthesized via Δ10 desaturation of palmitic acid (C_16_) followed by chain shortening, whereas the production of Z10-14:OAc involves the same FAD acting on myristic acid (C_14_) (Foster and Roelofs 1987; Hao et al. 2002; Albre et al. 2012). Our phylogenetic analysis of FAD genes shows that the *C. pomonella* genome contains several genes clustering with the tortricine Δ10 FADs, indicating that this biochemical activity may in theory still be carried out in olethreutines. Additional work will be necessary to elucidate the routes towards the biosynthesis of monounsaturated pheromone components with double bond at even positions.

Biological production of moth pheromones has been suggested as an environmentally-friendly alternative to conventional chemical synthesis. The concept is proven and large-scale production is on the horizon (Nešněrová et al. 2004; Hagström et al. 2013; Ding et al. 2014; Holkenbrink et al. 2020). Identification and characterization of the key enzymes underlying production of pheromone is a crucial step to identify the exogenous building blocks central to the engineering of plant and cell factories. In 2010, the estimated annual production of synthetic codlemone was 25 metric tons for the treatment by mating disruption of 200,000 ha of orchards worldwide (Witzgall et al. 2010). The identification and characterization of the desaturase involved in pheromone biosynthesis in *C. pomonella* is of particular interest as it may open up for biological production of codlemone in plant and cell factories by metabolic engineering (Löfstedt and Xia 2020) and represent an important step towards the biological production of pheromone for the integrated pest management of several economically important olethreutines.

## Materials and Methods

### Semiochemical Data Set

To compile a dataset of the semiochemicals used as sex attractants by tortricid moths we manually retrieved data from the Pherobase, a freely accessible database of pheromones and semiochemicals which compiles information from the literature and peer-reviewed publications (El-Sayed 2020). Altogether, we gathered data for 536 taxa belonging to the family Tortricidae. Specifically, we were interested in collecting information about the chemical structure of the bioactive fatty acyl derivatives, i.e. double bond position, number and length of the aliphatic chain. Following the database convention, we classified bioactive compounds as *pheromone* or *attractant*. The term pheromone denotes a component of the female sex attractant, identified from females, and with demonstrated biological activity; attractant characterizes those bioactive compounds found by male responses alone, such as by systematic field screening or by significant non-target capture in traps baited with pheromone lures. Although these compounds are not necessarily used in the natural communication system, they have in most cases a high probability of being pheromone components. Details are in Supplementary Table S1.

### Molecular Data Set

To infer the evolutionary relationship among the species represented in our trait dataset we used sequence data from 137 terminal taxa in our phylogenetic analysis, including 131 species of Tortricidae as ingroup. The outgroup contained six taxa, each a species representative of Cossidae, Galacticidae, Heliocosmidae, Lacturidae, Limacodidae and Sesiidae, respectively. Our data set is based largely on Regier et al (2012) and Fagua et al. (2017), augmented with a few additional Genbank accessions. Following the aforementioned authors, we used regions of five nuclear genes and one mitochondrial gene. The nuclear genes were: *carbamoyl-phosphate synthetase II* (CAD), *dopa decarboxylase* (DDC), *enolase* (Eno), *period* (PER), *wingless* (WG); the mitochondrial gene was the fragment of *cytochrome oxidase I* (COI) used for barcoding. Details are in Supplementary Table S2. For each genus, we used one representative species as the type species.Our main goal was to place the major transition in the pheromone communication system of Tortricidae and thus, to infer the phylogenetic relationships with good resolution at the subfamily and tribe level. We retrieved a total of 340 Genbank accessions (see Supplementary Table S2 for accession numbers).

### Phylogenetic Tree

We concatenated the molecular data corresponding to one taxon and obtained an alignment of the entire molecular data set using MAFFT (Katoh et al. 2002) as implemented in Geneious (Biomatters). The final alignment comprised 7591 sites. To define the best scheme of partitions using the MAFFT alignment, we used PartitionFinder v2.1.1 (Lanfear et al. 2017). We tested models of nucleotide substitution with partitions representing codon position and genes using *greedy* search (Lanfear et al. 2012), *AICC* model selection, and setting the phylogeny program to *phyml* (Guindon et al. 2010)(default value for all other settings). A 16-partitions scheme was selected and used to perform the parametric phylogenetic analysis in IQ-TREE 1.6.11 (Nguyen et al. 2015; Chernomor et al. 2016). The maximum likelihood analysis was performed using default settings and by calculating Shimodaira-Hasegawa-like approximate ratio test (SH-aLRT) support and ultrafast bootstrap support (UFBoot; (Hoang et al. 2018)) after 1000 replicates each. We used the R package *ggtree* (Yu et al. 2017) for visualizing and annotating the phylogenetic tree with associated semiochemical data.

### Insects, Tissue Collection, and RNA-Seq

Pupae of *C. pomonella* were purchased from Entomos AG (Switzerland) and were sexed upon arrival to our laboratory. Adult males and females emerged in separate jars in a climate chamber at 23±1°C under a 16h:8h light: dark cycle. On the third day after emergence, pheromone glands of female moths were dissected, immersed in TRIzol Reagent (Thermo Scientific), and stored at -80°C until RNA extraction. Two pools of 30 glands were collected 3 hours before and after the onset of the photophase, respectively. Total RNA was extracted according to the manufacturer’s instructions. RNA concentration and purity were initially assessed on a NanoDrop2000 (Thermo Fisher Scientific). Library preparation and paired-end Illumina sequencing (2×100bp) were performed by BGI (Hong-Kong, PRC). We obtained ∼66 million QC-passing reads per sample.

### Genome Annotation Improvements

To ensure that all FAD genes were correctly annotated in the C. pomonella genome we used our RNA-Seq data of pheromone gland to improve the annotation (cpom.ogs.v1.chr.gff3) which we obtained from InsectBase (http://www.insect-genome.com/cydia/). We performed low-quality base trimming and adaptor removal using cutadapt version 1.13 (Martin 2011) and aligned the trimmed read pairs against the genome using HISAT2 version 2.2.0 (Kim et al. 2019). The existing annotation was used to create a list of known splice sites using a python script distributed with HISAT2. We used StringTie version 2.1.3b (Pertea et al. 2015) with the *conservative* transcript assembly setting to improve the annotation and reconstruct a non-redundant set of transcripts observed in any of the RNA-Seq samples. We applied Trinotate version 3.2.1 (Bryant et al. 2017) to generate a functional annotation of the transcriptome data. We identified candidate FAD genes by searching for genes harboring the PFAM domain corresponding to a fatty-acid desaturase type 1 domain (PF00487) in their predicted protein sequences. We used the R package *KaryoploteR* (Gel and Serra 2017) to visualize the location of the FAD genes.

### Desaturase Phylogenetic Analysis

To investigate the evolutionary relationship of the *C. pomonella* FADs we carried out a phylogenetic analysis with other moth FAD proteins. Our sampling includes all the functional classes previously identified in moths (i.e. Δ5, Δ6, Δ9, Δ10, Δ11, Δ14) and representatives of Tortricidae as well as other moth families (see Supplementary Table S3 for list of taxa). Protein sequences for these enzymes were downloaded from GenBank. Protein sequences for *C. pomonella* were predicted from the genome using gffread (Pertea and Pertea 2020) with our improved annotation.

We aligned amino acid sequences using MAFFT version 7.427 (Katoh et al. 2002; Katoh and Standley 2013). We used the L-INS-i procedure and the BLOSUM30 matrix as scoring matrix. The final multiple sequence alignment contained 114 sequences with 805 amino-acid sites. The phylogenetic tree was constructed in IQ-TREE 1.6.11 (Nguyen et al. 2015). Automatic model search was performed using ModelFInder (Kalyaanamoorthy et al. 2017) with the search restricted to models including the WAG, LG and JTT substitution models. The maximum likelihood analysis was performed using default settings and by calculating Shimodaira-Hasegawa-like approximate ratio test (SH-aLRT) support and ultrafast bootstrap support (UFBoot; (Hoang et al. 2018)) after 1000 replicates each. We used the R package *ggtree* (Yu et al. 2017) for visualizing and annotating the phylogenetic tree, with midpoint rooting performed using the midpoint.root function implemented in the *phytools* package (Revell 2012).

### Transcript Abundance Estimation

In addition to our pheromone gland data sets, we retrieved 27 already published Illumina paired-end RNA-Seq data sets for *C. pomonella* available on the Sequence Read Archive (SRA) (see Supplementary Table S4 for accession numbers). These data correspond to various tissues or life-stages. Following QC, low-quality base trimming and adaptor removal with cutadapt, read pairs were aligned against the genome using HISAT2. The obtained alignment BAM files were used to estimate transcript abundance using StringTie together with our improved annotation. The abundance tables from StringTie were imported into R using the *tximport* package (Soneson et al. 2016), which was used to compute gene-level abundance estimates reported as FPKM. We used the R package *pheatmap* (Kolde 2019) to visualize the expression level of FADs genes.

### Cloning of Desaturases

First-strand cDNAs were synthesized from one microgram of total RNA using the SuperScript IV cDNA synthesis kit (Thermo Scientific). The cDNA products were diluted to 50 µL. PCR amplification (50 µL) was performed using 2.5 µL of cDNA as template with *attB*-flanked gene-specific primers (Supplementary Table S5) in a Veriti Thermo Cycler (Thermo Fisher Scientific) using Phusion polymerase master mix (Thermo Fisher Scientific). Cycling parameters were as follows: an initial denaturing step at 94°C for 5 min, 35 cycles at 96°C for 15 s, 55°C for 30 s, 72°C for 1 min followed by a final extension step at 72°C for 10 min. Specific PCR products were cloned into the pDONR221 vector (Thermo Fisher Scientific) in presence of BP clonase (Invitrogen). Plasmid DNAs were isolated according to standard protocols and recombinant plasmids were subjected to sequencing using universal M13 primers and the BigDye terminator cycle sequencing kit v1.1 (Thermo Fisher Scientific). Sequencing products were EDTA/ethanol-precipitated, dissolved in formamide and loaded for analysis on an in-house capillary 3130xl Genetic analyzer (Applied Biosystems).

### Pheromone Compounds and Corresponding Fatty Acid Precursors

All synthetic pheromone compounds used as references for identification came from our laboratory collection unless otherwise indicated. Pheromone compounds are referred to as short forms, e.g. (*E*8,*E*10)-8,10-dodecadienol is E8E10-12:OH, the corresponding acetate would be E8E10-12:OAc, corresponding acyl moiety (dodecadienoate) is E8E10-12:Acyl (although it may occur as CoA-derivative) and the methyl ester thereof is referred to as E8E10-12:Me. The dodecanoic methyl ester (methyl laurate, 12:Me) and tetradecanoic methyl ester (methyl myristate, 14:Me) were purchased from Larodan Fine Chemicals (Malmö, Sweden). The (*E*)-9-dodecenol (E9-12:OH) was purchased from Pherobank (Wageningen, The Netherlands) and then converted into the corresponding methyl ester (E9-12:Me). E8E10-12:OAc was purchased from Bedoukian (Danbury, CT, USA) and converted to the corresponding alcohol by hydrolysis using a 0.5 M solution of KOH in methanol. Fatty alcohols were oxidized to the corresponding acid with pyridinium dichromate in dimethylformamide as described by Bjostad and Roelofs (1984). The methyl esters were prepared as described under fatty acid analysis (see below).

### Heterologous Expression of Desaturases in Yeast

Plasmids containing the genes of interest were selected as entry clones and subcloned into the copper-inducible pYEX-CHT vector (Patel et al. 2003). Sequences of recombinant constructs were verified by Sanger sequencing. The pYEX-CHT recombinant expression vectors harboring *C. pomonella* FADs were introduced into the double deficient *elo1Δ*/*ole1*Δ strain (*MATa elo1::HIS3 ole1::LEU2 ade2 his3 leu2 ura3*) of the yeast *Saccharomyces cerevisiae*, which is defective in both desaturase and elongase functions (Schneiter et al. 2000) using the *S*.*c*. EasyComp Transformation kit (Thermo Fisher Scientific). For selection of uracil and leucine prototrophs, the transformed yeast was allowed to grow on SC plate containing 0.7 % YNB (w/o AA; with Ammonium sulfate) and a complete drop-out medium lacking uracil and leucine (Formedium, UK), 2% glucose, 1% tergitol (type Nonidet NP-40, Sigma), 0.01% adenine (Sigma) and containing 0.5 mM oleic acid (Sigma) as extra fatty acid source. After 4 days at 30°C, individual colonies were selected and used to inoculate 10 mL selective medium, which was grown at 30°C at 300 rpm in a shaking incubator for 48 h. Yeast cultures were diluted to an OD_600_ of 0.4 in 10 mL of fresh selective medium containing 2 mM CuSO_4_ for induction, with or without addition of a biosynthetic precursor. Yeast cells contain sufficient quantities of naturally-occurring palmitic acid, hence supplementation with 16:Me was typically not necessary. Lauric acid and myristic acid were added to the medium as methyl esters (12:Me and 14:Me). In subsequent assays the monounsaturated methyl esters E9-12:Me and Z9-12:Me were also supplemented. Each FAME precursor was prepared at a concentration of 100 mM in 96% ethanol and added to reach a final concentration of 0.5 mM in the culture medium. Yeasts were cultured in 30°C with Cu^2+^-induction. After 48 h yeast cells were harvested by centrifugation at 3000 rpm and the supernatant was discarded. The pellets were stored at –80°C until fatty acid analysis. Yeast expression experiments were conducted with three independent replicates.

### Heterologous Expression of Desaturases in Insect Cells

The expression construct for Cpo_CPRQ in the baculovirus expression vector system (BEVS) donor vector pDEST8_CPRQ was made by LR reaction. Recombinant bacmids were made according to instructions for the Bac-to-Bac^®^ Baculovirus expression system given by the manufacturer (Invitrogen) using DH10EMBacY (Geneva Biotech). Baculovirus generation was done using *Spodoptera frugiperda* Sf9 cells (Thermo Fisher Scientific), Ex-Cell 420 serum-free medium (Sigma) and baculoFECTIN II (OET). The virus was then amplified once to generate a P2 virus stock using Sf9 cells and Ex-Cell 420 medium. Viral titer in the P2 stock was determined using the BaculoQUANT all-in-one qPCR kit (OET) and found to be: 3 x 10^8^ pfu/mL for Cpo_CPRQ.

Insect cells lines Sf9 were diluted to 2 x 10^6^ cells/mL. Heterologous expression was performed in 20-ml cultures in Ex-Cell 420 medium and the cells infected at an MOI of 1. The cultures were incubated in 125 mL Erlenmeyer flasks (100 rpm, 27 °C), with fatty-acid methyl-ester substrates supplemented at a final concentration of 0.25 mM after one day. After three days, 7.5-mL samples were taken from the culture and centrifuged for 15 min at 4500*g* at 4° C. The pellets were stored at -80°C until fatty acid analysis. Sf9 expression experiments were conducted in three replicates.

### Fatty Acid Analysis

Total lipids from cell pellets were extracted using 3.75 mL of methanol/chloroform (2:1 v/v) in clear glass tubes. One mL of 0.15M acetic acid and 1.25 mL of water were added to wash the chloroform phase. Tubes were vortexed vigorously and centrifuged at 2000 rpm for 2 min. The bottom chloroform phase (∼ 1 m), which contains the total lipids, was transferred to a new glass tube. Fatty acid methyl esters (FAMEs) were produced from this total lipid extract. The solvent was evaporated to dryness under gentle nitrogen flow. One mL of sulfuric acid 2 % in methanol was added to the tube, vortexed vigorously, and incubated at 90°C for 1 hour. After incubation, 1 mL of water was added, mixed well, and 1 mL of hexane was used to extract the FAMEs.

The methyl ester samples were subjected to GC-MS analysis on a Hewlett Packard 6890 GC (Agilent) coupled to a mass selective detector HP 5973 (Agilent). The GC was equipped with an HP-INNOWax column (30 m × 0.25 mm × 0.25 µm; Agilent), and helium was used as carrier gas (average velocity: 33 cm/s). The MS was operated in electron impact mode (70eV), and the injector was configured in splitless mode at 220°C. The oven temperature was set to 80°C for 1 min, then increased at a rate of 10°C/min up to 210°C, followed by a hold at 210°C for 15 min, and then increased at a rate of 10°C/min up to 230°C followed by a hold at 230°C for 20 min. The methyl esters were identified based on mass spectra and retention times in comparison with those of our collection of synthetic standards.

Double-bond positions of monounsaturated compounds were confirmed by dimethyl disulfide (DMDS) derivatization (Buser et al. 1983), followed by GC-MS analysis. FAMEs (50 µL) were transferred to a new tube, 50 µL DMDS were added and the mixture was incubated at 40°C overnight in the presence of 5 µL of iodine (5% in diethyl ether) as catalyst. Hexane (200 µL) was added to the sample and the reaction was neutralized by addition of 50-100µL sodium thiosulfate (Na_2_S_2_O_3_; 5% in water). The organic phase was recovered and concentrated under a gentle nitrogen stream to a suitable volume (Liénard et al., 2008). DMDS adducts were analyzed on an Agilent 6890 GC system equipped with a HP-5MS capillary column (30 m × 0.25 mm × 0.25 µm; Agilent) coupled with an HP 5973 mass selective detector. The oven temperature was set at 80°C for 1 min, raised to 140°C at a rate of 20°C/min, then to 250°C at a rate of 4°C/min and held for 20 min (Wang et al., 2010).

Double-bond positions of di-unsaturated compounds with conjugated double-bonds were confirmed by derivatization with 4-methyl-1,2,4-triazoline-3,5-dione (MTAD) (Marques et al. 2004). MTAD adducts were prepared using 15 µL of the methyl ester extracts transferred into a glass vial. The solvent was evaporated under gentle nitrogen flow and 10 µL of dichloromethane (CH_2_Cl_2_) was added to dissolve the FAMEs. The resulting solution was treated with 10 µL of MTAD (Sigma; 2 mg/mL in CH_2_Cl_2_). The reaction was ran for 10 min at room temperature and 2 µL was subjected to GC-MS analysis on an Agilent 6890 GC system equipped with a HP-5MS capillary column (30 m × 0.25 mm × 0.25 µm; Agilent) coupled with an HP 5973 mass selective detector. The injector temperature was set to 300 °C, the oven was set to 100 °C and then increased by 15 °C/min up to 300 °C followed by a hold at 300 °C for 20 min.

## End Matter

## Author Contributions and Notes

J.-M.L, B.-J.D. and C.L. designed research; B.-J.D. collected tissue and processed samples for RNA-Seq; J.-M.L. performed analyses related to phylogenetics and bioinformatics; B.-J.D. performed functional assays and chemical analyses; J.-M.L. prepared the figures; J.-M.L. wrote the paper with input from B.-J.D. and C.L..

The authors declare no conflict of interest.

This article contains supporting information online.

## Acknowledgments

We thank the Lund Protein Production Platform (LP3) for technical support and advice on the expression in insect cell lines. The computations in this paper were run on the FASRC Cannon cluster supported by the FAS Division of Science Research Computing Group at Harvard University. This project has received funding from the European Union’s Horizon 2020 research and innovation programme (grant agreement n° 760798)(CL), the Swedish Foundation for Strategic Research (grant n° RBP 14–0037, Oil Crops for the Future) (CL), the Swedish Research Council (CL), Formas (CL) and the Royal Physiographic Society in Lund (JML).

## Appendix

### Supplementary figures

**Supplementary Figure 1:**
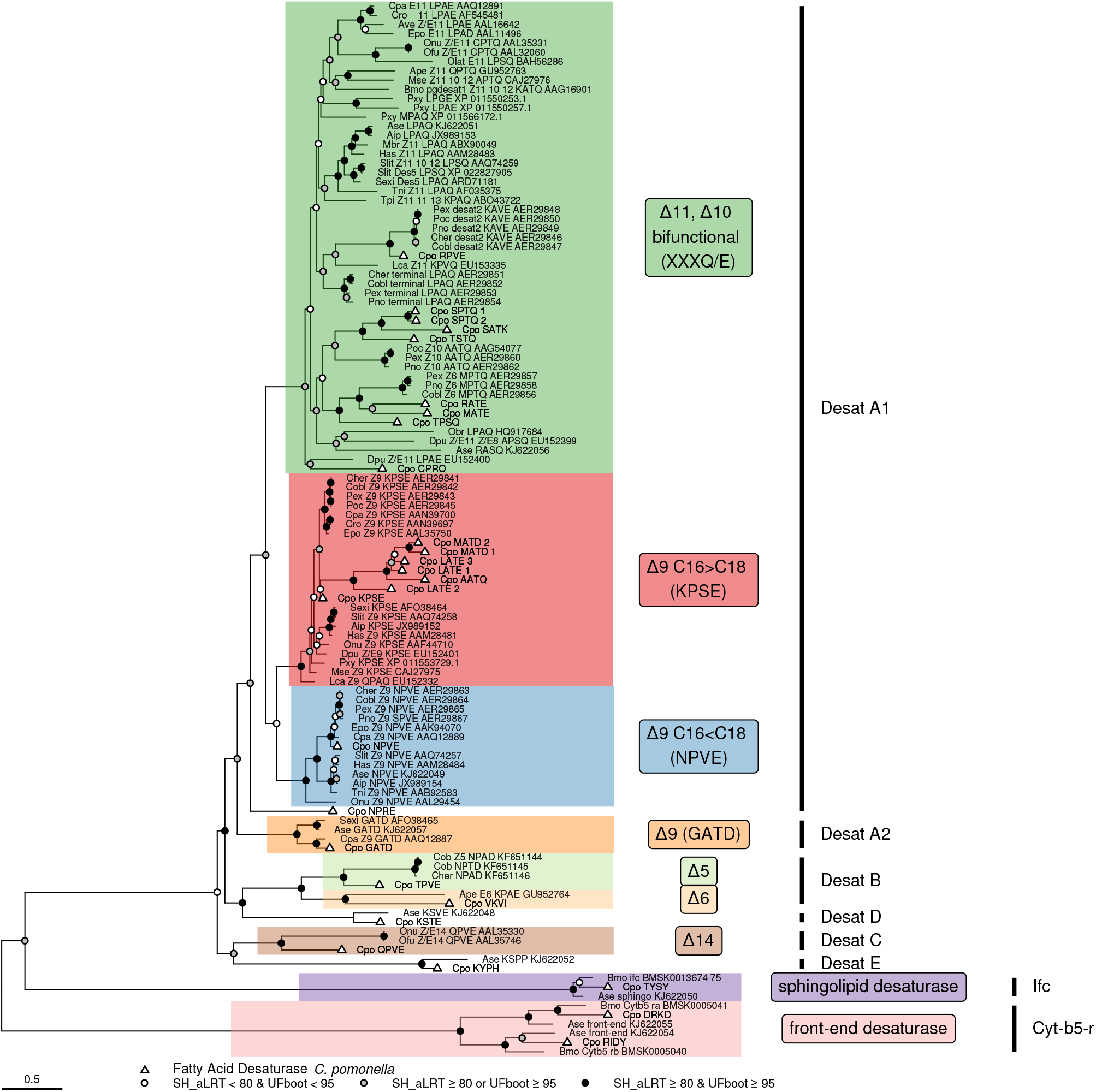
Phylogeny of lepidoptera FAD genes. Extended version of the maximum likelihood tree displayed in Fig 4. The tree was obtained for predicted amino acid sequence of 114 FAD genes (805 aligned positions) of 28 species, with branch support values calculated from 1000 replicates using the Shimodaira-Hasegawa-like approximate ratio test (SH_aLRT) and ultrafast bootstraping (UFboot). Support values for branches are indicated by colored circles, with color assigned based SH-aLRT and UFBoot supports with 80% and 95% as thresholds of branch selection for SH-aLRT and UFBoot supports, respectively. The major constituent six subfamilies of First Desaturase (A1 to E) and two subfamilies of Front-End (Cyt-b5-r) and Sphingolipid Desaturases (Ifc), respectively, are indicated following the nomenclature proposed by Helmkampf et al. (2015). For First Desaturases, the different shades correspond to the indicated putative biochemical activities and consensus signature motif (if any). Triangles indicate sequences from *C. pomonella*. The scale bar represents 0.5 substitutions per amino acid position. Species are indicated by three- or four-letter prefixes (see Supplementary Table S3 for details). Biochemical activities (or signature motif) are indicated after the abbreviated species name, followed by the accession number.

**Supplementary Figure 2:**
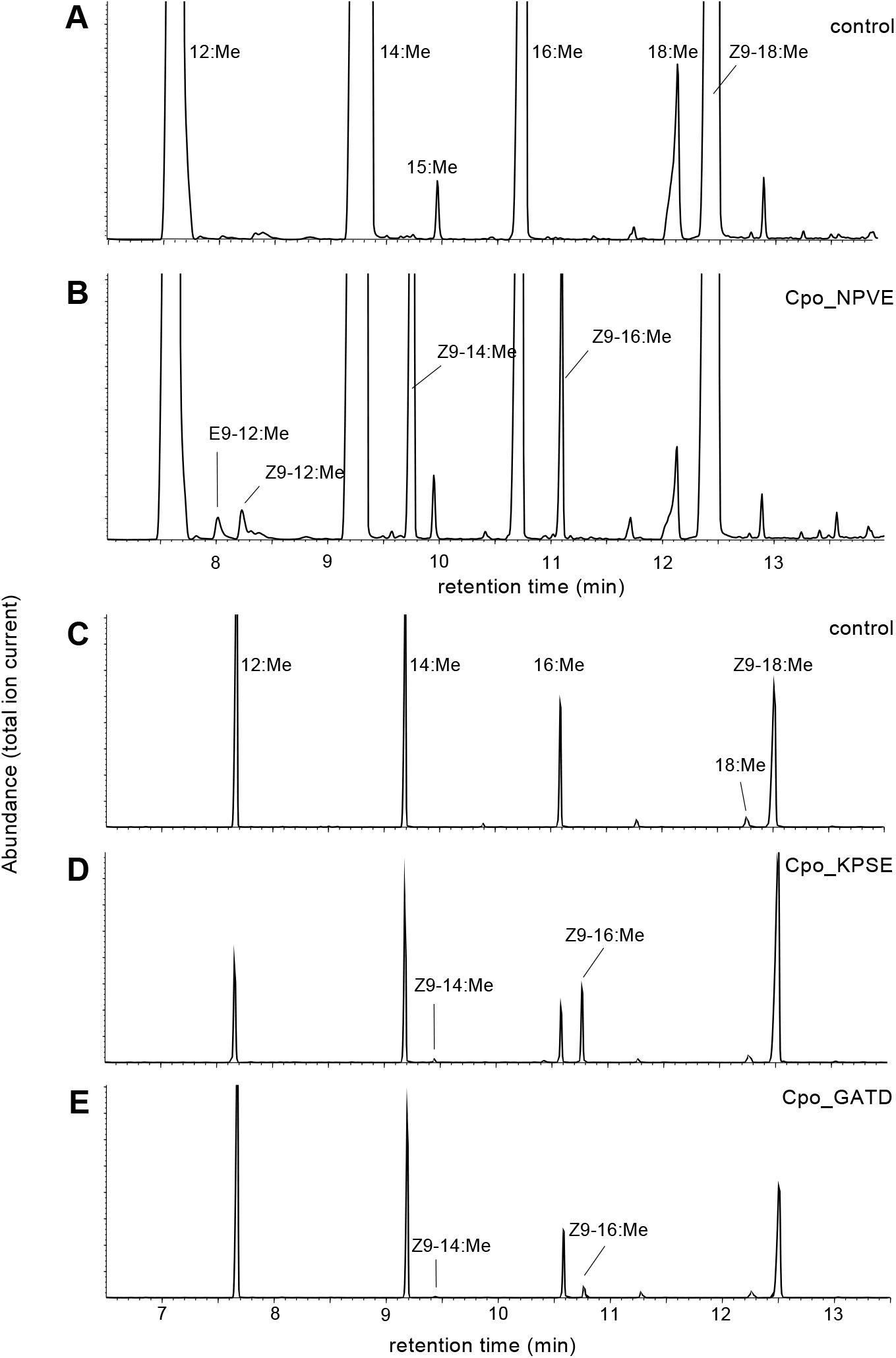
Functional characterization of desaturase activity of First-desaturases. Total ion chromatograms of fatty acid methyl ester (FAME) products of Cu^2+^-induced *ole1 elo1 S. cerevisae* yeast supplemented with saturated acyl precursors and transformed with (A &C) empty expression vector (control), (B) pYEX-CHT-Cpo_NPVE, (D) pYEX-CHT-Cpo_KPSE, and (E) pYEX-CHT-Cpo_GATD.

